# Widespread variation in heat tolerance of the coral *Acropora hyacinthus* spanning variable thermal regimes across Palau

**DOI:** 10.1101/2020.04.26.062661

**Authors:** Brendan Cornwell, Katrina Hounchell, Nia Walker, Yimnang Golbuu, Victor Nestor, Stephen R. Palumbi

## Abstract

Climate change is poised to dramatically change ecosystem composition and productivity, leading scientists to consider the best approaches to fostering population resilience and diversity in the face of these changes. Here we present results of a large-scale experimental assessment of bleaching resistance, a critical trait for coral population persistence as oceans warm, in 293 colonies of the coral *Acropora hyacinthus* across 39 reefs in Palau. We find bleaching resistant individuals originate significantly more often from warmer reefs, although they inhabit almost every reef regardless of temperature at low frequency. High levels of variation *within* reefs, where colonies experience similar temperatures, suggests that bleaching resistance is not solely due to phenotypic plasticity, but also involves adaptive alleles and host-symbiont interactions. To the extent that it is heritable, bleaching resistance could be used in promoting nursery growth, habitat restoration, or breeding, while employing large numbers of resistant colonies to preserve genetic variation.

## INTRODUCTION

Climate change is increasingly shifting species ranges, altering ecosystem dynamics, and generating strong selection differentials in wild populations (Chen et al. 2011; Logan et al. 2014; MacLean and Beisinger 2017; Wiens 2016; Parmesan & Yohe 2003; Chen et al. 2011; Lenoir & Svenning 2015). Against this backdrop, there is an increased focus on characterizing the adaptive mechanisms that will increase resilience to climate stressors in natural communities, particularly on the role that heritable genetic variation and selection will play in allowing populations to adapt to future conditions (King et al. 2018; Walsworth et al. 2019). For instance, numerous studies have uncovered variation in heat resistance within species of fish, lizards, corals, marine plants, and a host of aquatic and terrestrial species (King et al. 2018; Eliason et al. 2011; Herrando-Pérez et al. 2019; Somero 2010; Oliver and Palumbi 2011; Stuart-Smith et al. 2017). This variation, particularly differences in environmental tolerance among individuals or populations, provides the raw material on which selection will act to increase the resilience of populations facing future climate change (Logan et al. 2014; Nyamukondiwa et al. 2018).

The practical importance of the potential adaptive response of a species is rooted in the possibility that populations already harboring stress tolerant individuals might be used in restoration projects or in assisted evolution efforts to enhance the resilience of vulnerable populations (Aitken and Whitlock 2013; Mascia & Mills 2017; Seddon et al. 2014). For instance, forest restoration projects have tended to use seed from local source trees because they were long thought to represent the population most adapted to local conditions. However, the onset of rapid climate change now requires considering the use of seed stock from distant populations that currently experience a set of conditions resembling those that the restored population of interest is likely to encounter in the future (Prober et al. 2015), effectively resulting in assisted gene flow across the landscape. Norberg et al. (2012) and Walsworth et al. (2019) showed that models that included evolutionary responses to climate change stress could lead to much higher levels of stable persistence than ecological models without evolution. The key parameter in models that predict population persistence is increased heritable phenotypic variation, which allows a population to evolve quickly enough to match changing environmental conditions. Identifying source populations with such phenotypic variance could become especially critical for foundation species such as forest trees, sea grasses, and corals, in order to maintain current levels of productivity, function and diversity of ecosystems into the future (Franks et al. 2014; Hodgins and Moore 2016, Morikawa and Palumbi 2019).

Yet, there are serious limitations to using non-local populations in restoration. Long distance transplantation for conservation or restoration can come with unintended impacts such as introducing diseases, invasive species, or devastating pests (Ricciardi and Simberloff 2009). Furthermore, the loss of locally adapted phenotypes through gene swamping (Menkhorst et al. 2016; Weeks et al 2011; Sullivan, Nowak, & Kwiatkowski 2014; Plein et al. 2015) can have deleterious consequences if selectively advantageous alleles for traits not directly linked to surviving climate change such as salinity, disease or water quality tolerance (e.g. Jin et al. 2015) are reduced in frequency or eliminated. In such circumstances, propagating individuals using a limited gene pool can reduce overall fitness if assisted evolution and/or gene flow programs designed to mitigate the effects of climate change fail to evaluate the suitability of source individuals for traits other than heat resistance (NASEM 2019a). The best strategy to minimize these problems is to use a large population of locally collected stress-resistant individuals, which highlights a fundamental gap that undermines the implementation of these conservation efforts: the spatial scales over which managers can find a large number of stress tolerant individuals are unknown for most species.

A key starting point for these studies is a census of adaptive variation for environmental stress-resistance in wild populations (Bush et al. 2016). These surveys provide a baseline measurement of how well a species can adapt to different conditions. They can also provide guidance to locations where stress-resistance is most commonly found for a species and give an indication of how frequently stress resistance arises in a population. In an early study, Meuli and Shirley (1937) used a common garden to test seedlings of the green ash tree and drew a simple map of resistance to drought across the US Midwest in order to facilitate seed collection and planting of drought resistant ash groves. Several studies have generated environmental stress resistance maps for a wide number of terrestrial plants and animals (e.g. Schmidt et al. 2005; Sorenesen et al. 2009; Weldon et al. 2018) as the first step to discovering the mechanisms of stress resistance or making use of these traits in replanted or restored ecosystems. Similarly, environmental heterogeneity across the seascape will certainly produce non-random distributions of stress-tolerant individuals and characterizing their distribution is fundamental for predicting how these populations will respond to climate change.

As with many terrestrial and marine ecosystems, coral reefs are highly vulnerable to climate change and other anthropogenic stressors (e.g. Hughes et al. 2003; Frieler et al. 2013). In recent years, global reef-building coral populations have declined as a result of bleaching events, disease and storms linked to warming oceans (Heron et al. 2016; Hughes et al. 2017). Nevertheless, corals living in naturally occurring regions and across long-distance latitudinal gradients can be differentially adapted to high temperatures (Dixon et al 2015; Berkelmans and Willis 1999, Sawall et al. 2015). In addition, research at very small spatial scales has shown that corals growing in warm microclimates are more resistant to bleaching (Palumbi et al. 2014) and that individual coral colonies within a population can exhibit variable heat tolerance (Oliver and Palumbi 2011; Barshis et al. 2013; Morikawa and Palumbi 2019). Here, we extend work on small scale variation in heat tolerance by conducting standardized heat stress tests on 293 table top corals of the species complex *Acropora hyacinthus* across 39 reefs in the northern and southern lagoons of Palau. We included shallow patch reefs likely to experience highly variable, warm conditions, and deeper fore reefs which more consistently experience cooler oceanic water temperatures. By mapping and testing individual corals we generated a fine-scale heat tolerance map. The results show a wide distribution of heat tolerant colonies, concentrated in warmer patch reefs, but which are also present on cooler fore reefs and deeper habitats. Moreover, substantial phenotypic variance for heat tolerance is present in 34 of 39 reef populations. The large inventory of heat tolerant colonies across the archipelago, could provide ample, local stocks of corals for natural evolution of heat tolerance, and is the foundation for future work on assisted gene flow, nursery construction or selective breeding.

## MATERIALS AND METHODS

### Experimental Protocol

#### Site Selection and Temperature Measurements

We selected 10 coral colonies per reef from 39 reefs distributed around the Palauan archipelago in 2017. In order to fully capture the range of potential phenotypic responses to warming, we chose reefs that encompass a wide range of naturally occurring water temperatures around the island. Reefs also ranged in size from approximately 5,000 m^2^ to >100,000 m^2^, and in depth ranging from ca. 1-8 m at low tide. We selected 15 reefs in the northern lagoon of Babeldoab island, where surface waters are regularly replaced by cooler oceanic waters flushing in with the tides (Skirving et al. 2005), and 15 reefs in the southern lagoon where water is retained for longer periods and has the potential to warm to higher temperatures. We selected patch reefs in grouped clusters that typically contained at least 3 reefs that were near one another. Additionally, we selected five fore reefs near each of the northern and southern patch reef regions (although we could not re-sample one of the northern fore reefs to run the bleaching assay), for a total of 39 reefs.

To identify 10 colonies on each reef, we haphazardly swam transects from the exterior to the interior of each patch reef to capture the full range of abiotic and biotic conditions at each location. We selected fore reef colonies by swimming parallel to the reef crest, just inside the edge of each reef. We tagged each colony with a unique numeric identifier (PA1-400), and recorded its location using a handheld GPS (Garmin, Kansas). Additionally, to characterize the thermal environment, we deployed a HOBO temperature logger (OnSet Computing, Massachusetts) recording at ten-minute intervals to all odd-numbered colonies (five per reef). Temperature data were collected at all locations from November 8, 2017 to July 20, 2018 (see Supplementary File 1A for temperature records).

#### Fragment Collection and Stress Tank Protocol

We experimentally stressed fragments from each of the 400 corals that we tagged and monitored for this study. We clipped medium sized fragments (approximately 8cm width) from the edge of each colony using garden clippers, loosely packaged the fragments in bubble wrap and stored them in a cooler to be transported to the Palau International Coral Reef Center (PICRC). Upon return, fragments were placed in a flowing seawater system at ambient temperatures where they recovered from transport overnight. The following day, we clipped the larger fragments into five smaller fragments and placed them in our experimental warming system.

Each of the 10 L experimental tanks was independently controlled using a 300W heater and 2 chillers (Nova Tec, Maryland), and had fresh seawater inflow at all times (ca. one volume change in 2-3 hours) in addition to an aquarium pump (∼ 280 L hour^-1^) to increase flow around the fragments. Tanks were illuminated on a 12 hour light:dark cycle using LED light fixtures (ca. 22-66 μmol m^-2^s^-1^). Three experimental treatments ramped a fragment from each colony to temperatures of 34°C, 34.5°C, or 35°C, with two additional control tanks that did not ramp but were maintained at 30°C for the duration of the experiment. Each experiment consisted of two ramping cycles over the course of two days. Ramps began at 10:00 A.M. and increased temperature for three hours until they reached their target temperature at 1:00 P.M. Heating lasted for three hours (until 4:00 P.M.) at which point chillers cooled each tank back to 30°C, typically within 30 minutes. When tanks were not ramping, they maintained a constant temperature of 30°C. We repeated this cycle over two days, after which corals were held at 30°C overnight before preservation. We preserved coral tissue by removing tissue from each fragment using an airbrush loaded with seawater. We then centrifuged the tissue for five minutes at 5,000 g, removed the supernatant and resuspended the slurry in 2 ml of RNALater. We stored these samples at 4°C for approximately 24 hours, after which time we transferred them to −20°C until shipping to Hopkins Marine Station.

We visually evaluated coral fragments at 8:00 A.M. each day of the experiment (one observation before each ramp and a final observation the day following the second ramp for a total of three observations). We visually assessed coral fragments using a five point scale that consisted of the following categories: 1 – no bleaching, 2 – visually discolored, 3 – moderately discolored, clearly bleaching, 4 – severely discolored, nearly complete bleaching, 5 – no color, completely bleached. We evaluated each fragment using two scorers who were required to agree on a score for each fragment during each assessment. We also took photographs of each tank immediately following the visual bleaching score assessment for confirmation in cases where the visual score disagreed with symbiont quantification by flow cytometry (described below).

#### Quantification of Symbiont Load

In order to more precisely quantify the symbiont density in each fragment after experimental heating, we used a Guava EasyCyte HT (Millepore, Massachusetts) flow cytometer to count symbiont cells. To prepare each sample, we centrifuged 500 μl of each cell suspension sample plus 500 μl filtered seawater at 15,000 g for 5 minutes, removed the supernatant and resuspended each pellet in 300 μl 0.01% SDS solution. We homogenized each sample using a PowerGen rotostat (Fisher Thermosciences, Massachusetts) set at the highest setting for 5 seconds. We then needle sheared each sample through a 25-gauge needle 10 times to ensure cells were not clumped together. Each sample was then diluted by 1:200 in 0.01% SDS and run in triplicate on the flow cytometer. We gated sample counts on the forward scatter channel, counting all events exceeding 10^2^ fluorescent units. Additionally, we counted all events that exceeded 10^4^ fluorescence units on the 690 nm detector (which can detect the autofluorescence of chlorophyll) as a symbiont count. After subtracting events we detected in the negative control (0.01% SDS), we calculated the proportion of symbionts as the total number of symbiont events divided by the total number of events above the forward scatter gate. For a subset of samples, we also quantified the total protein content of the homogenized (undiluted) sample using a Pierce BCA protein assay (Thermo Scientific, Massachusetts) as per protocols in Krediet et al. (2015). We measured the absorbance of these samples in triplicate on a Tecan Plate Reader at 562 nm. We note that while cell count values may not represent absolute numbers of cells within tested colonies (e.g. due to cell fragmentation during tissue airbrushing and preservation, or Guava cell counting error), values do confidently represent relative differences between samples.

## ANALYSIS

### Geographic Temperature Variation

Previous work found that oceanic waters routinely flush through the interior of northern fore reefs during tidal cycles (Skirving et al. 2005), which might make these environments colder than southern interior waters. Similarly, we also expected fore reef environments that are exposed to cooler upwelling conditions to have fewer extreme temperature events than protected patch reef environments. The thermal profiles that we recovered from loggers deployed on the same reef were highly correlated, we therefore used the average thermal profile for each reef for these analyses. After trimming the temperature measurements to begin and end at the same time points, we tested these predictions by comparing the mean number of time events above 31°C in patch reefs compared to fore reefs using a one-tailed t-test. Other temperature thresholds resulted in similar rankings. We then conducted a comparison between northern and southern patch reefs using a one-tailed test with the prediction that northern reefs should experience fewer high temperature events than southern reefs.

### Geographic Patterns of Bleaching Resistance

We examined whether the rate of bleaching resistance over the course of the two-day experiment was dependent on geographic region (northern vs. southern reefs). First, we calculated the average change in visual bleaching score (VBS) from day zero (no heat stress) to day one (one cycle of heat stress) and from day zero to day two (two cycles of heat stress). We then used a t-test to detect differences in bleaching rates between those two regions and within regions across days of the experiment.

We normalized the reduction in symbiont density for each temperature treatment by dividing the density of symbionts remaining after heat stress by the density of the control treatments. We then used these standardized values to assess the degree to which corals bleached during heat stress. We determined the performance of each coral as the fraction of the average proportion of symbionts that were present after the heat treatments compared to the average of the two control fragments. We also estimated the temperature at which each colony lost 50% of its original symbionts. To do this we assumed a linear decline in symbiont density from the initial symbiont proportions in the control treatment to the final proportions in the three temperature treatments. From this linear model we then estimated the temperature treatment that would be necessary to elucidate the loss of half of the initial symbiont population

We based heat resistance on three different criteria. The first was the proportion of symbionts in heated branches versus control branches of the same colony, termed symbiont retention. The second criterion was the change in Visual Bleaching Score during heating for branches compared to their controls. Colonies were deemed heat resistant by each of these criteria if they were in the upper 25%. The third criterion was final Visual Bleaching Score of 4 or less at 35°C. (For several colonies we used VBS at 34.5 when there were no VBS data at 35°C. Because VBS was 0.7 units higher on average at 34.5°C than 35°C, we used a VBS score of 3 or less at 34.5°C). There were 105 corals (36%) that fit this third criterion.

We tested if bleaching resistant colonies were evenly distributed across reefs in Palau. Without incorporating other environmental factors, we tested whether the proportion of bleaching resistant individuals was the same across the five geographic regions that contained the 39 reefs we sampled (Eastern, South Central, Western South Lagoon, Fore reefs, and Northern reefs). We used a chi-square test to test the hypothesis that bleaching-resistant colonies are represented in equal proportions across those regions.

Bleaching resistant colonies should reside in warmer environments if selection results in a subset of individuals that are adapted to that environment or individuals can acclimate to local conditions. One prediction if the thermal environment selects for heat resistant colonies, or results in acclimation to increased temperature, is that bleaching resistant colonies should reside in warmer environments. We therefore assessed whether bleaching responses were correlated with the number of extreme temperature events on each reef. Because data loggers deployed to each reef were highly correlated (see above), we used the average number of time points above 31°C for all loggers on a reef to represent their exposure to thermally challenging conditions. We also conducted these analyses using increasing thresholds of 32°C, 33°C and 34°C, none of which substantially changed the outcome (not shown).

To test the relation of local heating to heat resistance, we used a fully factorial linear model where the average fraction of the symbiont population that was lost in the heated treatments (measured by flow cytometry) was the dependent variable and the number of extreme temperature events on a reef, the area of the reef (excluding fore reef sites), and the average depth of the colonies on each reef were the independent variables.

Finally, we characterized the proportion of each reef that is comprised of heat resistant and sensitive corals (defined as any coral meeting 2 out of 3 criteria described above, a total of 74 bleaching resistant and 42 bleaching prone corals were included).

## RESULTS

### Cell count and visual score data

We recorded control and heated symbiont density data from 361 colonies across 39 reefs. We removed 35 colonies from the dataset for which we had only one control value (n=30) or only one heat treatment value (n=5). We also removed an additional 33 colonies for which the heated branches showed anomalously higher density of symbionts than did the controls (ratio range 4.4 to 1.2). Most of these colonies (n=20) showed low symbiont proportions in the controls (<0.05). These changes left us with a filtered cell count data set of 293 colonies (Supplementary File 1B).

Initial symbiont proportions measured by flow cytometry were highly heterogeneous in corals across the archipelago. Mean symbiont proportions in the control treatments for the 293 colonies ranged from 0.69% to 26.12% at the end of the experiment (Fig. 2A). However, symbiont proportions do not correlate with temperature extremes (see above) across reefs (r^2^=0.003679 p=0.1792), nor do they differ statistically between northern (0.0981) and southern (0.0888) reefs (t = −1.7293, df = 280.09, p-value = 0.0849).

We measured Visual Bleaching Scores (VBS) for 1608 branches from these colonies after one and two days of heating at 30, 34, 34.5 and 35°C. A comparison between VBS and the proportion of symbionts measured with flow cytometry shows high correlation (Fig. 1). Coral fragments scored to have no bleaching (VBS=1) on average show 12% of their cells to contain symbionts whereas totally bleached corals (VBS=5) show 3%. However, there is high variance in symbiont proportions within each VBS category, reflecting both the variation among non-heated colonies (Fig. 1) and the wide variation in bleaching states that are lumped into each visual bleaching category.

**Figure 1.**
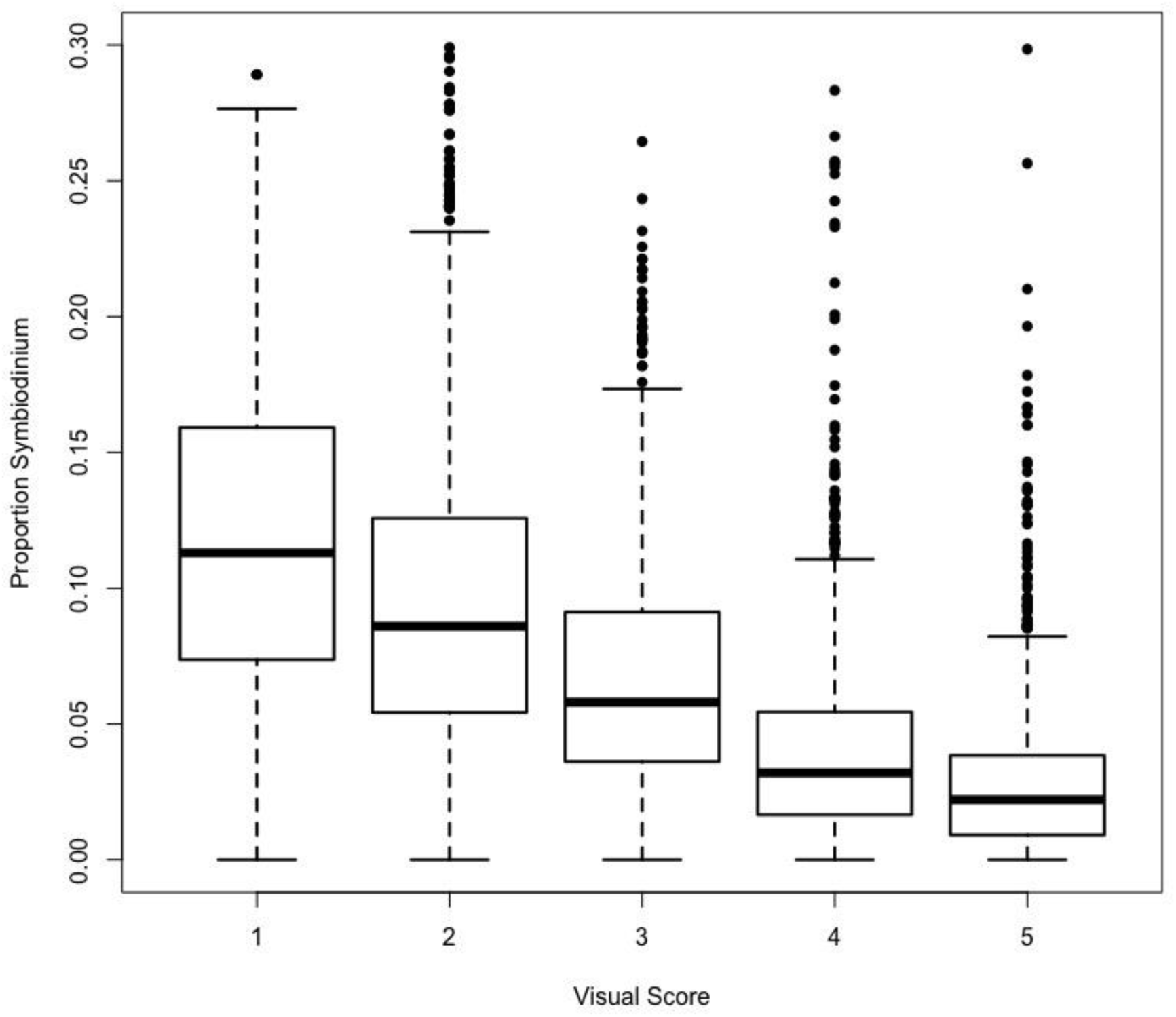
Range of symbiont cell concentration by visual bleaching score category. Symbiont cell proportions are highly correlated with visual bleaching score, though there is high variance particularly among colonies with low bleaching scores. Note: measurements are of all colonies from the experimental heat treatments as well as the control treatments. Accompanying source data are available as Figure 1 – Source Data.

### Symbiont densities after standardized experimental bleaching

The different rates of bleaching we observed in this study were largely due to individual reactions to heat stress on the first day of the experiment followed by much smaller reductions on day 2. First, baseline measurements across all experiments on day zero show that on average, northern corals started the experiment with higher symbiont densities (mean VBS = 1.8) than their southern counterparts (mean VBS 2.0, t=2.093, df=283.37, p=0.0372). From those initial densities, the decline in VBS in northern locations between day zero and day one was approximately 45.2%, while the scores of individuals from southern locations declined approximately 39.3% by the end of day one (t=2.889, df=282.91, p=0.00416). Northern and southern populations continued to bleach between day one and day two, but at nearly identical rates. Northern individuals lost an additional 6.9% of their initial symbiont populations as measured by the VBS, while southern individuals lost 6.4%. The symbiont population densities in both populations were significantly lower on day two compared to day one, ending at total losses of 52.1% and 45.7%, respectively (t=3.379, df=282.85, p=0.000830).

On average colonies retained 53% of their original symbionts after heating for two daily cycles as measured by flow cytometry. However, different colonies showed wide variation in symbiont retention after standardized heating, ranging from 100% retention to less than 5% (Fig. 2A). We saw >90% retention of symbionts after heating in 31 corals from 18 reefs. By contrast, 45 corals from 24 reefs retained less than 25% of their symbionts under the same experimental conditions. Bleaching-resistant corals (in the top 25% of retention) averaged 75% symbiont retention whereas bleaching-prone corals (bottom 25%) averaged 37% retention (Fig. 2A).

**Figure 2.**
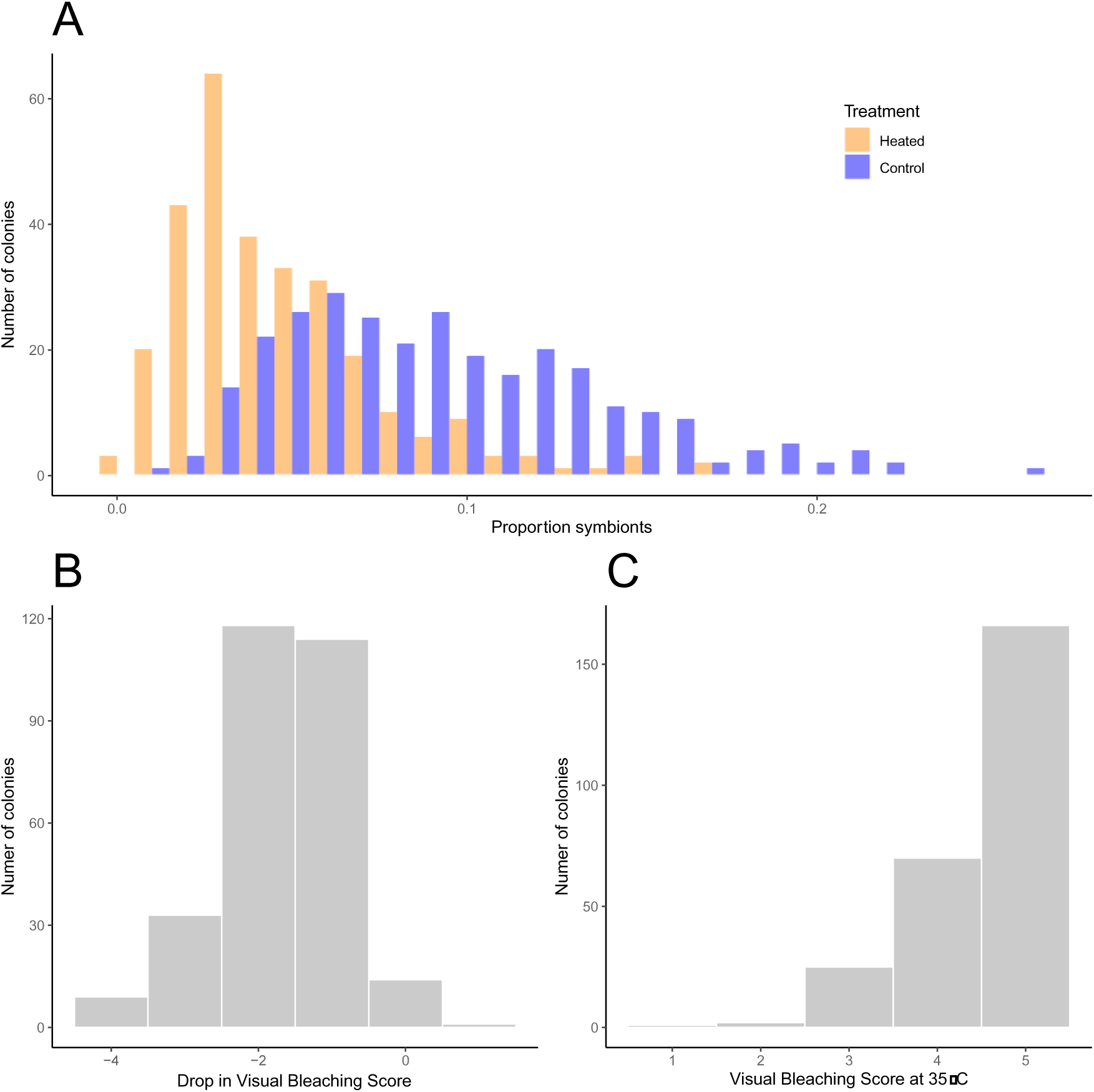
Variation in heat tolerance using three stress test response criteria. A: Mean proportion of symbionts in tissue cells from corals before and after heating. B: Decreases in visual bleaching scores after heating. C: final visual bleaching scores after heating at 35°C. See Figure 2 – supplemental 1 for further comparison of the change in visual bleaching scores compared to changes in symbiont proportions after heat treatment. Accompanying source data are available as Figure 2A – Source Data and Figure 2B and C – Source Data.

To test if our characterization of heat resistance based on the retention of symbionts was robust, we also scored heat resistance based on the drop in Visual Bleaching Score compared to controls. Increasing VBS were correlated with decreasing symbiont proportions (r^2^=0.3054, p<2 x 10^-16^, Fig. 2 – supplemental 1). The difference (delta) in VBS and symbiont cell counts between control treatments and heated treatments was also correlated but did not explain a large portion of the variance (r^2^=0.1287, p<2×10^-16^, Fig. 2 – supplemental 1). Because these analyses suggest that VBS and cell counts capture slightly different aspects of bleaching, we incorporated metrics of bleaching resistance based on the visual bleaching scores alone. Colonies ranged from no change to a VBS shift of four categories, with a change of one category or less (e.g. category 1 to category 2) occurring in 42 corals across 19 reefs. By contrast 22 colonies showed a VBS change of 3-4 (Fig. 2B).

Last, we also used the visual bleaching scores for colonies after two days at the highest temperature, 35°C, to find colonies with the highest heat resistance. Most colonies (171) showed total bleaching at this temperature but about 24% showed severe bleaching and 10% showed only moderate bleaching or less (Fig. 2C).

We used these three measures of heat resistance (proportion symbiont retained, drop in visual bleaching score, and absolute visual bleaching score at the highest temperature) to characterize each colony we tested. We considered a colony to be highly heat resistant for a criterion if it was in the upper 25% −34% of colonies. For symbiont retention, this included all colonies that retained more than 73% of their symbionts after heating (average 89.4%). For visual bleaching scores, our highly resistant category included all colonies that dropped less than one visual bleaching value (85 colonies or 30%), and all colonies that showed less than total bleaching at the highest temperature stress (97 colonies or 34%). There are 29 colonies that are bleaching resistant based on all three criteria, and another 45 that are highly resistant on the basis of two of three criteria (Supplementary File 1B).

Corals that meet 2-3 heat resistance criteria start with slightly but insignificantly fewer symbiont cells compared to corals that meet none of those criteria (9% vs 9.7%, t=1.4576, df=148.19, p=0.1471), but show much higher levels of symbionts after heating (average 6.3% vs 3.3%, p=3.743×10^-9^), resulting in retention rates of 75% versus 37% (Table 2). Likewise, Visual Bleaching Scores were the same in controls for both categories (2.4 vs 2.3, t = 1.8309, df = 141.66, p = 0.0692), but differed strongly when corals meeting 2-3 criteria versus those meeting none were heated (VBS of 3.7 versus 5.0 at 35°C, Table 2).

**Table 1.** Temperature extremes by geographic region. Minimum and maximum number of temperature events above 31°C in the five geographic regions in this study.

|  | Western South<br>Lagoon | Eastern | Forereef | South<br>Central | North<br>Lagoon |
| --- | --- | --- | --- | --- | --- |
| Minimum | 2395 | 1547 | 151 | 1204 | 205 |
| Maximum | 3204 | 1845 | 1526 | 2652 | 1933 |

**Table 2:** Comparison of bleaching resistant and prone individuals. Average results of bleaching experiments for corals that meet 2-3 heat resistance criteria versus those that meet none.

| Category | Proportion<br>symbionts<br>in controls | Proportion<br>symbionts<br>at 34 °C | Proportion<br>symbionts<br>at 34.5 °C | Proportion<br>symbionts<br>at 35 °C | Ave.<br>proportion<br>symbionts in<br>heated<br>colonies | VBS Day 2<br>controls | VBS Day 2<br>34°C | VBS Day 2<br>34.5°C | VBS Day 2<br>35°C | Retention | VBS heated<br>minus cntrl |
| --- | --- | --- | --- | --- | --- | --- | --- | --- | --- | --- | --- |
| Meets 2-3 criteria | 0.090 | 0.073 | 0.065 | 0.047 | 0.063 | 2.368 | 2.608 | 3.108 | 3.721 | 0.753 | 0.749 |
| Meets no criteria | 0.097 | 0.041 | 0.030 | 0.021 | 0.033 | 2.328 | 3.942 | 4.241 | 5.000 | 0.369 | 2.094 |

### The geography of heat resistant corals

Corals that are heat resistant on the basis of at least two of our three criteria are found at many locations: at least one heat resistant colony (out of the 10 we surveyed) inhabited 31 of 39 reefs (Fig. 3 upper). However, heat resistant corals tend to be more common on warmer patch reefs on the western and eastern edges of the southern lagoon (red and green bars in Fig. 3; Chi Square, X^2^=12.476, df=4, p=0.01414). For example, 14 of the 29 corals that met all three heat resistant criteria were from the eastern (Reefs 7 and 9) or the western patch reefs in the southern lagoon (PR 22-27, Fig. 3 lower). By contrast, fewer bleaching resistant colonies are found on patch reefs in the northern lagoon (4 of 29, Fig. 3), or on the south-central patch reefs (2 colonies on Reefs 17, 18, 19 Fig. 3). Fore reef locations overall show about the expected number (17 observed, 13 expected) but these occurred only on four reefs, three of which were on the western edge of the northern barrier reef (Reefs 60, 61, 62, purple bars). Overall, the number of colonies that met all three criteria was higher in southern reefs (17 observed) than in northern reefs (12 observed, t=3.7104, df=261.69 p=2.53 x 10^-4^).

**Figure 3:**
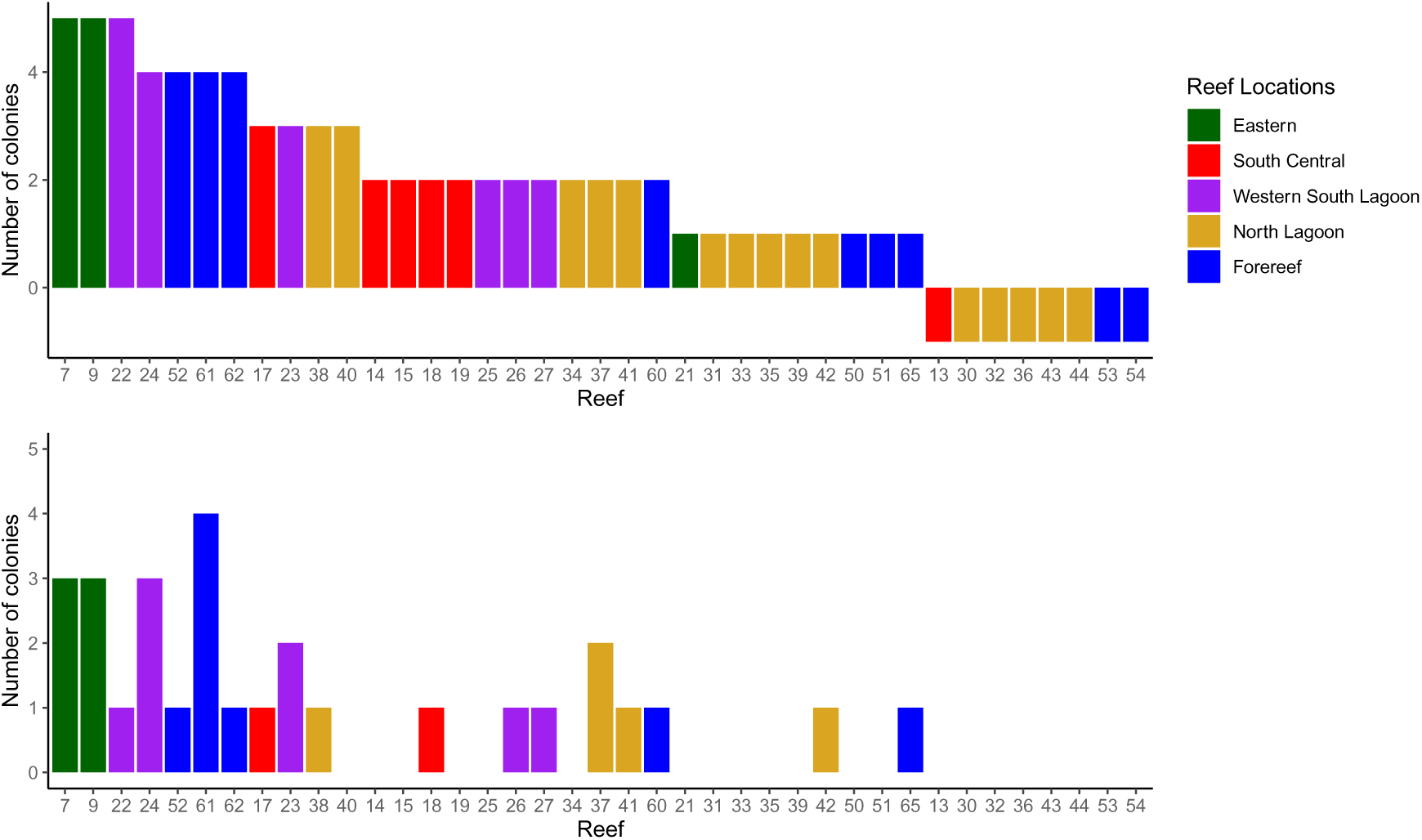
Location and prevalence of heat resistant colonies by region. Upper: Location of corals that show top 25% scores in two of three heat resistance criteria. Lower: Location of corals that show top 25% scores in three of three heat resistance criteria. Accompanying source data are available as Figure 3 – Source Data.

### Geographic Patterns in Temperature

We compared temperature data from 77 corals on 32 reefs, recorded from November 8, 2017 to July 20, 2018 (Supplementary File 1A). Overall, fore reef locations and patch reefs in Palau’s northern lagoon had lower temperatures and fewer high temperature extremes than did patch reefs in the southern lagoon. Fore reef sites showed an average of 983 temperature events above 31°C (out of 60,960 periods each lasting 10 minutes, or about 3.2% of time periods) compared to patch reef environments which have an average of 1784 (5.8% of time periods, t=4.1709, df=57.659, p=5.165 x 10^-5^). However, patch reefs were highly divergent depending on location: patch reefs in the North lagoon had, on average, 865 extreme temperature events, about the same as fore reefs, compared to 2460 in the southern sites (t=8.129, df=56.975, p=2.105 x 10^-11^). The total range for the number of extreme temperature events for the entire study across reefs was 151-3204.

Extreme temperature events on reefs tended to cluster regionally (Fig. 4A and Table 2). For example, a set of patch reefs on the north-western edge of the southern lagoon (PR22-27) shows some of the highest temperatures in our study, where 2400-3200 time periods exceed 31°C (Fig. 4A). By contrast, fore reef locations were more uniformly cool (151-1526 temperature records above 31°C; Supplementary File 1A). By contrast, some reefs located near one another showed dramatically different thermal regimes. For example, reefs in the south-central part of the southern lagoon are located within 5 km of each other (e.g. PR 17, 18, 19, Supplementary File 1A) but showed a range of values, spanning 1204-2652 events over 31°.

**Figure 4.**
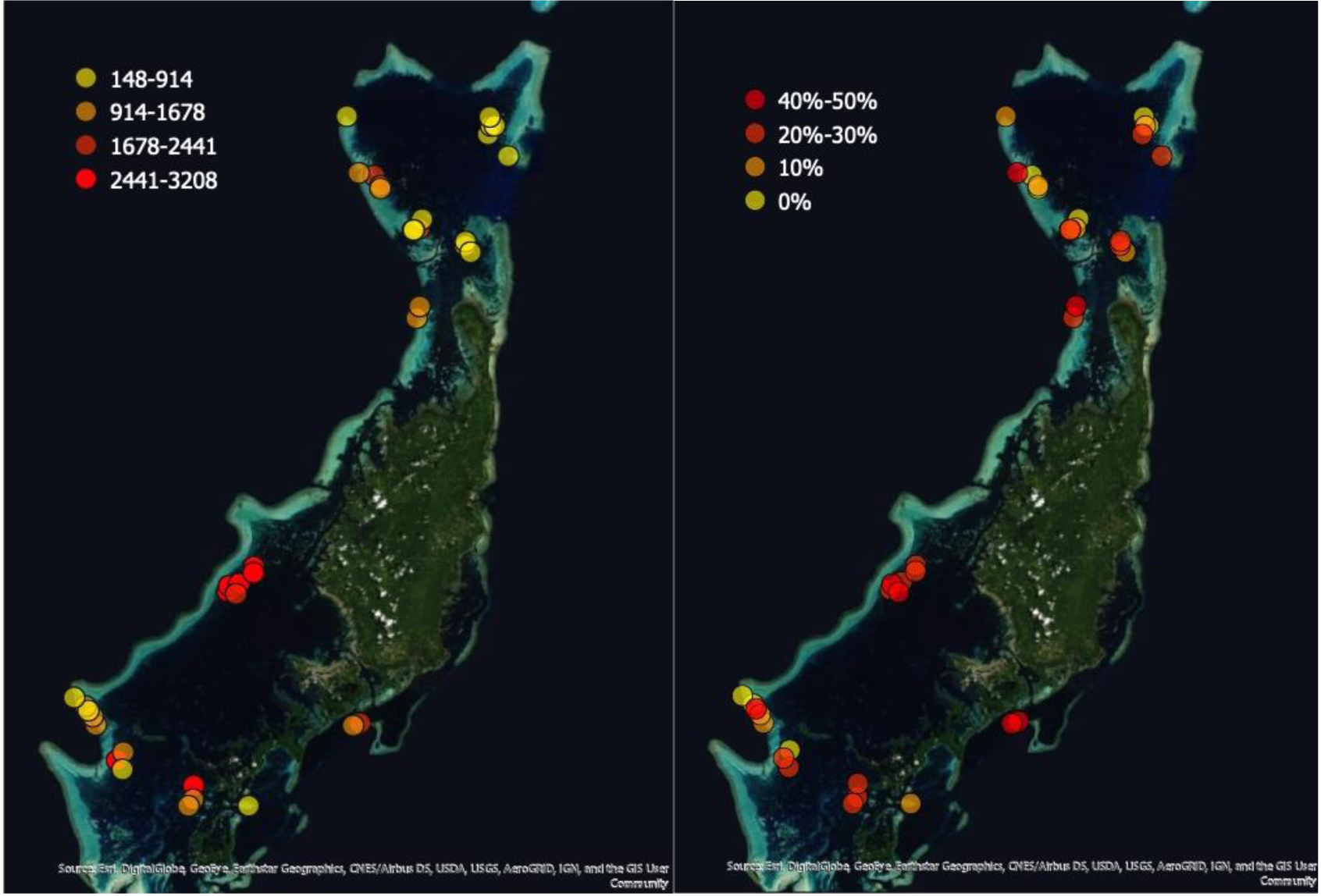
Distribution of bleaching resistance and reefs experiencing warm conditions. *Left*: Fraction of time periods exceeding 31°C on each reef. Patch reefs on the eastern and western edges of the southern lagoon tend to have the most common warm periods. Mean number of event periods exceeding 31°C for each of the 39 reefs in this study. See Supplementary File 1A for details. The total number of time periods compared was 60,960. Differently colored dots represent exposure up to 1.5%, 3.0%, 4.5% and 6.0% of the time at 31°C. *Right*: The distribution and frequency of bleaching resistant colonies across the Palauan archipelago. See figure 4 – supplemental 1 for the results of the linear model comparing the number of bleaching resistant individuals on a reef to the number of extreme temperature events that reef experienced.

Although the variance is high, bleaching resistant corals generally were found on warmer reefs: they experienced an average of 1,868 warm events compared to bleaching susceptible corals (those that met none of our heat tolerance conditions) which experienced 1,316 warm events on average (t=3.859, df=114.52, p= 1.89 x 10^-3^). The number of corals that met 2 of 3 heat resistance criteria on a reef is also correlated with the number of warm temperature events that reefs experienced during this study (Fig. 4 - Supp.1). However, even most cooler reefs in our study had one or two heat resistant colonies, suggesting that factors other than temperature contribute to the distribution of bleaching resistance.

### Variance in heat tolerance

Our analysis concentrates on comparing mean bleaching values, symbiont retention, and visual bleaching scores for corals across standard experiments. However, patterns of phenotypic variance are also important because this variance, if heritable, allows selection to operate on the population.

Variance in bleaching was not consistent across the reefs we surveyed – on average the variance in the temperature at which colonies showed 50% retention of symbionts was 0.6 (mean 34.5°C, Supplementary File 1A). High variance occurred on reefs PR09, and PR22-27, which also had some of the highest mean symbiont retention values (Supplementary File 1A): these values were driven by a few colonies with exceptionally high levels of retention. By contrast, the far northern reefs (PR40-44) showed low heat resistance and low variance. Overall, all but a few reefs had substantial variation in heat tolerance among corals. Variance in heat resistance was positively correlated across reefs with mean symbiont retention (R^2^=0.49, p<0.05), reef temperature (R^2^=0.30, p<0.05), and the fraction of corals meeting 2 of 3 criteria (R^2^=0.22, p<0.05, one tailed) but not reef depth or size.

Much of the variance in heat tolerance is distributed among colonies, with smaller amounts distributed among reefs or reef regions. In particular, geographic region (North and South) explained ∼0% of the variance, whereas reef cluster explained 5.12%, the individual reef level explained 5.76% (mixed effects model of symbiont retention after bleaching as the dependent variable and reef temperature as the independent variable with geographic region and reef clusters as fixed effects). The large bulk of variance (ca. 89%) was partitioned at the individual colony level.

### Bleaching intensity and symbiont load

Bleaching intensity and initial symbiont load were related in the colonies we tested for this study. Before our bleaching tests, top corals had almost 20% less symbiont load than did bottom corals (0.0877 vs 0.107, t=2.3224, df = 113.58, p= 0.0220). Top reefs were split between southern lagoon locations where corals tended to start with high symbiont loads, and northern fore reef locations where many corals started with low symbiont proportions (Fig. 5). For example, nine of the 30 corals from reefs FR60-62 had initial symbiont loads averaging 3.7% symbionts per coral cell. After bleaching these same corals had 2.9% symbionts, but these pale corals are recorded as having high retention (77%) because they started out with few symbionts. Symbiont load was not correlated with the number of extreme temperature events each individual colony experienced over the course of the study and a linear model explained very little variance (F=3.12, df=1, 220, p=0.0787, r^2^=0.0095).

**Figure 5.**
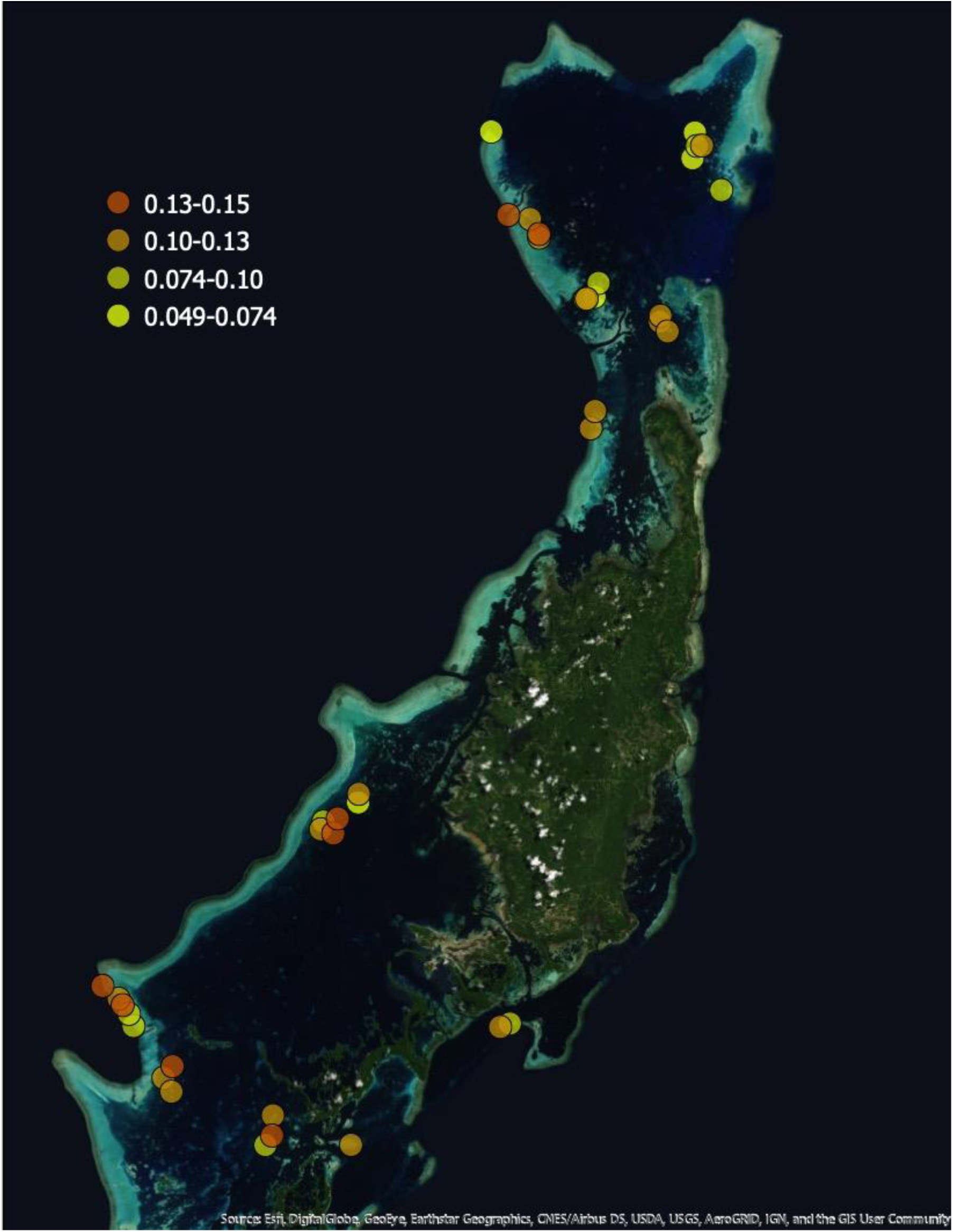
Mean symbiont load of *A. hyacinthus* colonies across Palau. Average proportion of symbionts in control treatment nubbins at the end of the experiment across the 39 reefs in this study. See figure 5 – supplemental 1 for the results of a linear model comparing initial symbiont load with symbiont retention under heat stress.

### Traits of reefs with high heat tolerance

Reefs varied in location, size, depth, number of events over 31, variance in temperature, and surrounding sea surface temperature. The number of events over 31°C and the variance in temperature both showed substantial correlation with the mean coral heat tolerance. We found no evidence that any of the abiotic factors we measured for each reef predicted the symbiont retention and heat tolerance of corals on that reef. We ran a fully factorial linear model where individual colony symbiont retention was the dependent variable with the following predictor variables: reef area, number of time points above 31**°**C, and average depth of colonies. None of these variables alone or interacting with another were significant predictors of symbiont retention.

## DISCUSSION

Across 39 reefs that we sampled in Palau, we found wide variation in bleaching susceptibility resulting from heat stress of the table top coral *A. hyacinthus*. In a simple two-day standardized heat stress experiment, colonies ranged from retaining virtually all of their original symbiont load at 35°C (ca. 5°C above ambient temperatures) to less than 30% at this temperature (Fig. 2). Overall, 10% of the 293 colonies we sampled met all three of our criteria for high bleaching resistance, and another 15% met two of three criteria; if this fraction is representative of the species as a whole, then the absolute number of bleaching resistant colonies across the Palau archipelago is large and they are distributed across a wide range of environments.

If heritable, the natural variation we identified in these populations is the foundation on which selection can act due to differential mortality from high temperatures. Given the current pace of climate change, selective mortality of bleaching-prone individuals due to heat stress is likely to be a strong force shaping the evolution of coral populations. This is already evident on the Great Barrier Reef, where coral populations may already be experiencing selection for bleaching resistance (Hughes et al. 2018). However, simulation studies conducted by Walsworth and colleagues show that conservation strategies that preserve habitat diversity can capitalize on the adaptive capacity of a population to mitigate the impacts of warming oceans (Walsworth et al. 2019). Likewise, Bay et al. (2017) showed that natural levels of genetic diversity in cool water coral populations could allow evolution of heat resistance as water temperatures warmed under mild CO_2_ emission scenarios. Consequently, the distribution and abundance of heat resistant colonies is a key feature of future reef resilience, and possible interventions to enhance it.

### Local geography of heat resistance

Bleaching resistant colonies are associated with warmer reefs in Palau. For example, the western-most patch reefs in the southern lagoon showed the highest number of high-temperature events during our survey (average of 2941 events, 4.8% of recorded time periods) and showed the largest number of colonies meeting 2-3 of our heat resistance criteria (18 of 60). This is consistent with previous studies, which have shown that coral populations harboring bleaching resistant individuals tend to undergo more short-term stress due to aerial exposure (Schoepf et al. 2015), live in shallow back reefs (Oliver and Palumbi 2011), or inhabit intrinsically warm ocean basins such as the Persian Gulf (Coles and Riegl 2013). In Palau, bleaching resistant coral communities tend to inhabit near-shore bays, where water temperatures are warmer and some of the highest bleaching impacts occur in northern reefs that are consistently at cooler temperatures than those in the South (van Woesik et al. 2012). However, patch reef corals did not bleach less than fore reef corals during natural bleaching events (van Woesik et al. 2012). This might be attributable to higher temperatures on patch reefs during bleaching periods. However, results in van Woesik et al. (2012) were highly variable among coral genera. For example, *Acropora* on fore reefs bleached less than their patch reef counterparts and *Pocillopora* showed the opposite, although these differences were not large. It should be noted that small-scale variation in thermal regimes (*e.g.* PR 17 and to a lesser extent FR 60 and 61 are particularly warm relative to nearby reefs) could be contributing to bleaching resistance, making environmental designations based on reef morphology alone difficult to interpret in the context of identifying locations likely to harbor bleaching resistant corals. Overall, whether coral assemblages are well adapted to their local temperature regimes is an area for continued research.

Nevertheless, we found substantial numbers of heat resistant colonies on reefs that experienced very few extreme temperature events. For example, cooler patch reefs in Palau’s northern lagoon had 17 heat resistant colonies. Although this was a low percentage of the colonies we surveyed, these individuals added substantially to our inventory of heat resistant colonies. This is also true of shallow fore reef locations, which harbored 17 heat resistant colonies in our final data set (Fig. 3 and Fig. 4B).

These results show that bleaching resistance due to heat stress is not always restricted to the warmest environments. Although we found patch reefs that had a high prevalence of bleaching resistant colonies were also associated with a large number of extreme temperature events, notably the western edge of the southern lagoon, fore reef settings are the most extensive habitat for *A. hyacinthus* (Veron and Stafford-Smith 2000), and they may paradoxically house the majority of heat resistant colonies in this species in Palau. Finding these colonies will be more difficult because they only occur at a frequency of 18% in our survey. However, high census population sizes on fore reefs means that they could contain the highest numbers of bleaching resistant individuals.

### Symbiont and environmental contributions to heat resistance

Overall, the mechanisms that create a mosaic pattern of bleaching resistance among colonies on a reef can fall into several broad categories with very different underlying causes: symbiont type, environmental history, and host genetics.

In particular, the symbionts themselves could be contributing significantly to variable resistance among coral colonies. Coral holobiont heat tolerance is known to vary with the genotypes and physiologies of the single-celled dinoflagellate endosymbionts that carry out photosynthesis within host tissues (e.g. Sogin et al. 2017). In particular, *Durusdinium* symbionts tend to confer higher heat resistance to the holobiont (Berkelmans et a 2006l; LaJeunesse et al. 2009). In a common garden experiment in American Samoa, for example, differences in heat resistance were correlated just as highly with differences in symbiont as they were with coral host genetics (Morikawa and Palumbi, 2019). Fore reef and patch reef corals in Palau largely host *Cladocopium* sp. symbionts with small populations of *Durusdinium* sp. (Quigley et al. 2014). To date we have identified no strong differences in symbiont composition using genus specific PCR patterns, but further tests of generic and species-level genetic differences are needed to fully understand the varied roles of coral and symbiont in our results.

A second well-known feature of coral heat tolerance is that warm water conditions induce higher heat tolerance (Middlebrook et al. 2008; Palumbi et al. 2014; Seneca and Palumbi 2015; Kenkel and Matz 2016; Ainsworth et al. 2015; Bay and Palumbi 2017). In our data, higher heat resistance of corals on warmer reefs could be partially due to this effect. However, marked variance among corals within the same reef suggests that the environment is not the sole driver of the patterns we documented. For example, on moderately warm patch reef PR09, four corals meet all three criteria of high heat resistance (PA11, PA12, PA14, PA15). By contrast, two colonies within 100 meters meet none of our criteria and are among the ten least heat resistant corals in our study (PA19 and PA20). Although microclimates within reefs can differ, temperature data from PA11, PA15, and PA19 show high consistency, with R^2^ values between colonies of 0.86 to 0.91. These data suggest that these colonies are experiencing very similar thermal regimes despite their differences in heat tolerance.

Strong bouts of selective mortality should alter phenotypes in populations where traits that mitigate environmental stress are heritable. Selection has been shown to change allele frequencies in mussels settling in estuaries, barnacles and snails in the high intertidal, abalone affected by strong harmful algal blooms, and fish under size selective fishing pressure (Koehn 1980, Schmidt and Rand 2001, Johannsen et al. 1995, de Wit et al. 2015, Therkildsen et al. 2019). In instances where shifts in abiotic conditions occur across a landscape (e.g. due to changing latitude or altitudes) multiple selectively advantageous alleles at a locus can be maintained across the geographic range of a species, typically in the form of clines in allele frequencies correlated with environmental conditions (Lewontin 1974). In general, evolutionary theory predicts that these polymorphisms are maintained by tradeoffs – where alleles that are adaptive in one habitat are detrimental in another (Levene 1953, Maynard Smith 1974).

In the case of corals, the adaptive value of heat tolerance is clear (see bleaching records in Rose et al. 2018). However, bleaching resistance should sweep to fixation in the population unless some countervailing force reduces the fitness of individuals enjoying its benefits. Paradoxically, the search for tradeoffs between heat resistance and other traits has yielded few convincing examples. For example, in a reciprocal transplant experiment, Bay and Palumbi (2017) found a consistent but only marginally significant tendency for corals that survived the best in the warm environment to grow more slowly in the cool environment. Likewise, Mieog et al. (2009) found mild and sporadic tradeoffs between growth and survival in *Acropora millepora*.

Alternatively, it could be that directional selection is acting on the population but has yet to fix beneficial alleles. Anthropogenic heat stress is becoming more common in tropical marine systems but when viewed through the lens of long-lived corals with generation times on the order of decades, has only selected for heat tolerance on a handful of *A. hyacinthus* generations. Northern patch reefs experienced strong coral death after the 1998 beaching event (Golbuu 2007) but have experienced few heat episodes since then. Although selection differentials are large when widespread bleaching events occur (Hughes et al. 2018), an insufficient number of generations coupled with so few years of severe heat stress across the reefs of Palau may have yet to yield fixation of adaptively beneficial alleles.

### Using heat resistant colonies for further coral reef resilience

This study used simple, standard heat stress tests to identify bleaching resistant corals across the Palauan archipelago and has revealed that bleaching resistant colonies inhabit a wide range of reef habitats and thermal environments. The map of heat resistance (Fig. 4A, supplemental video) shows both the wide geographic extent and frequency of heat resistance in the archipelago of Palau. This distribution implies that populations where warming has not yet caused widespread bleaching are likely to contain bleaching resistant individuals. If this variation results from intrinsic genetic variability, demonstrated using common garden studies, then it could form the basis for adaptation to climate change in the future, and could be used for three different conservation improvements.

The first is protection – reefs such as those on the western edge of the southern lagoon could be protected from overwhelming damage from avoidable human impact such as overfishing, pollution, or development. The second is use of these colonies to increase local heat resistance. Simulations have shown that an infusion of heat resistant genes can increase population adaptability and preserve census sizes under some future emission scenarios (Bay and Palumbi 2017). This kind of infusion might be orchestrated by building local coral nurseries with heat resistant colonies. Recently, Morikawa and Palumbi (2019) have shown that choosing heat resistant colonies of three species results in healthy nursery populations that resist bleaching 2-3 times better than nurseries built from heat sensitive colonies from the same reefs. This represents a possible target for resilience engineering of coral populations: using local heat resistant colonies to inject heat resistant alleles into local populations. A third possible conservation strategy is breeding of heat resistant corals to create offspring with even higher heat resistance (e.g. Kenkel et al. 2015). An important caveat to the last two strategies is that assisted gene flow and migration are only feasible if individuals can inter-breed; the widespread and interspersed distribution of cryptic *A. hyacinthus* species (Ladner and Palumbi 2012; Sheets et al. 2018) with varying heat tolerances (Rose et al. 2018; Gold and Palumbi 2018) requires additional confirmation that transplanted individuals are from the same species as the local population. The results from the present study suggest that conservation activities within relatively small spatial scales (10’s to 100’s of km) can capitalize on the local availability of heat resistant colonies (and the local diversity of cryptic species) to buffer populations from the impact of warming water temperatures.

Conservation strategies are beginning to consider the movement and selective breeding needed to mitigate the impacts of climate change in corals (NASEM 2019b) and other systems (Aitken and Bemmels 2016). These interventions have the potential to move pathogens or deleterious alleles and can potentially disrupt coevolved interactions between host and symbiont populations (Kranabetter et al. 2012). Using individuals sourced locally for local restoration efforts would reduce the risk of introducing pathogens from distant locations into an immunologically naïve population, which is a risk that has long been known when transporting individuals for other purposes such as commercial fisheries (Sindermann 1992). Furthermore, alleles that are adaptively beneficial in the local environments of Palau, but not involved in bleaching resistance, will continue to afford higher fitness to individuals originating from reefs within the archipelago.

### Conclusions

This study represents the first to comprehensively test individual colonies from a reef-building coral species for their heat resistance capacity across a large number of island-wide reefs with varying thermal regimes. Our results have demonstrated there are significant differences in heat resistance between individuals, and that resistant colonies are found more often on warmer reefs. However, we found corals with high levels of heat resistance even in cooler environments. There are likely additional adaptive and environmental factors that collectively explain the observed patterns, and future research should focus on disentangling the roles heritable and non-heritable variation play in the bleaching resistance of this species.

Finally, we have shown that identification of bleaching resistance does not necessarily require specialized equipment. Our standard test setup requires a temperature controller with reasonable accuracy, a heater, and a source of cooler fresh seawater. Our basic data collection protocol in the field uses a simple five-point visual bleaching scale. Although there was considerable variation in symbiont density for each visual bleaching score level, the agreement between the visual scores and symbiont proportions measured using flow cytometry, when combined with the ease of collecting visual scores relative to flow cytometry measurements makes this approach especially effective at rapidly and inexpensively assaying coral populations. Visual bleaching scores miss some important details (*e.g.* that symbiont load is negatively correlated with bleaching severity), however, the ease of documenting strong differences in bleaching results with visual bleaching scores – severe versus visible bleaching for example – opens up this tool to wider adoption. This will be of use not only to scientists, but importantly also to local managers with limited project budgets and manpower.

## ACKNOWLEDGEMENTS

The authors would like to acknowledge the staff and boat operators of the Palau International Coral Reef Center, as well as Julien Ueda, Mica Chapuis, Bowen Jiang, Callan Hoskins, Collin Hyatt, and Mehr Kumar for their assistance in the field.

## SUPPLEMENTAL FILES

Supplemental methods and results

Figure 2 – supplement 1

Figure 4 – supplement 1

Figure 5 – supplement 1

Supplementary File 1 (A and B)

## SOURCE DATA FILES

Figure 1 – Source Data

Figure 2 – supplemental 1 – Source Data

Figure 2A – Source Data

Figure 2 B and C – Source Data

Figure 3 – Source Data

Figure 4 – supplemental 1 – Source Data

Figure 5 – supplemental 1 – Source Data

## RICH MEDIA FILES

Depiction of the distribution of heat resistant colonies of *A. hyacinthus* across the Palauan archipelago.

---

**Figure 2 – figure supplement 1.**
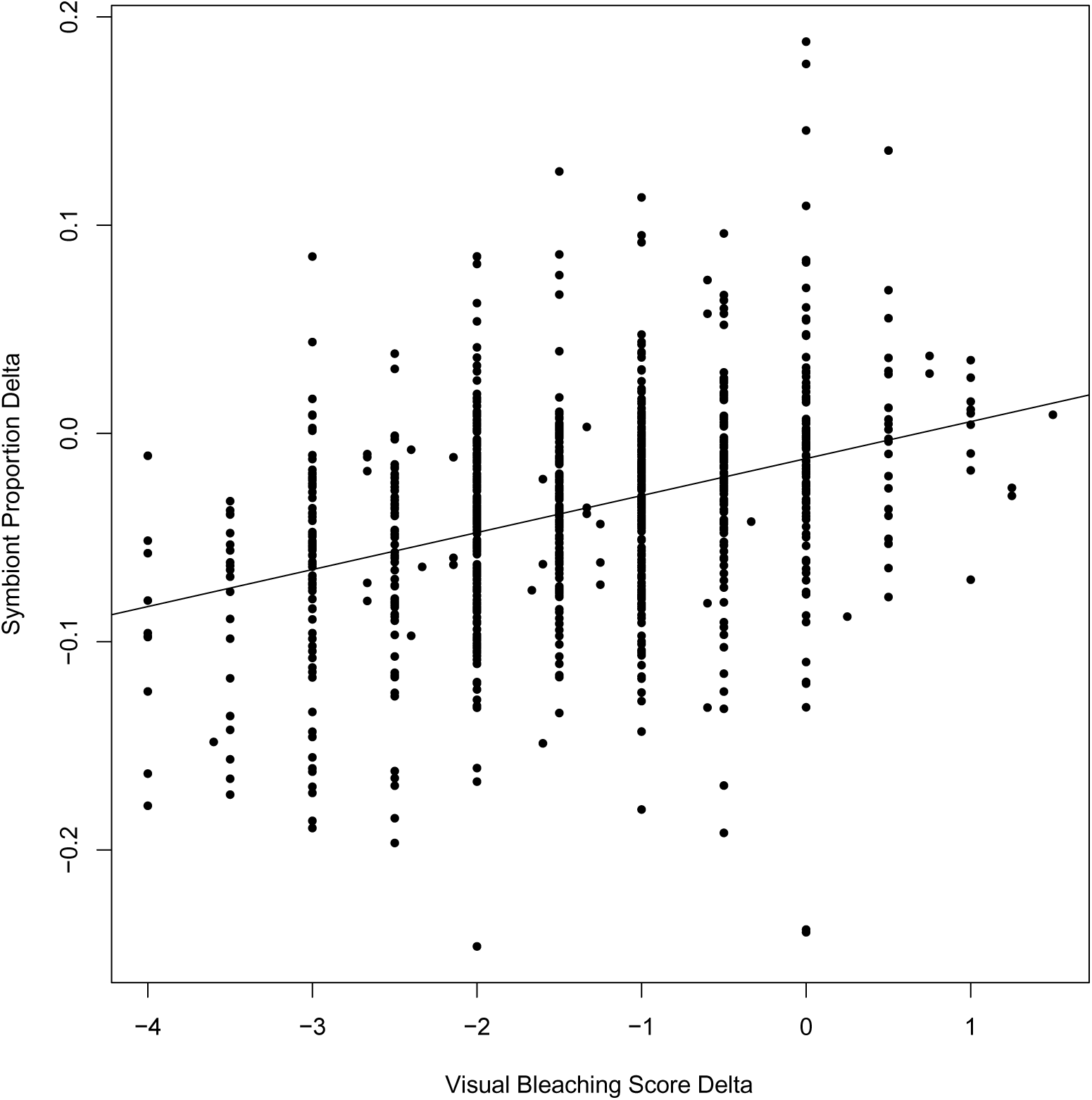
Change in symbiont proportion compared to change in visual bleaching scores when comparing control treatments to heat treatments across all colonies in this study. Each point represents a difference between a heat treatment and control. Thus, a single colony is represented by three points, one for each heat treatment relative to the average control values. R^2^=0.1287, p<2×10^-16^. Accompanying source data are available as Figure 2 – figure supplement 1 – Source Data.

**Figure 4 – figure supplement 1:**
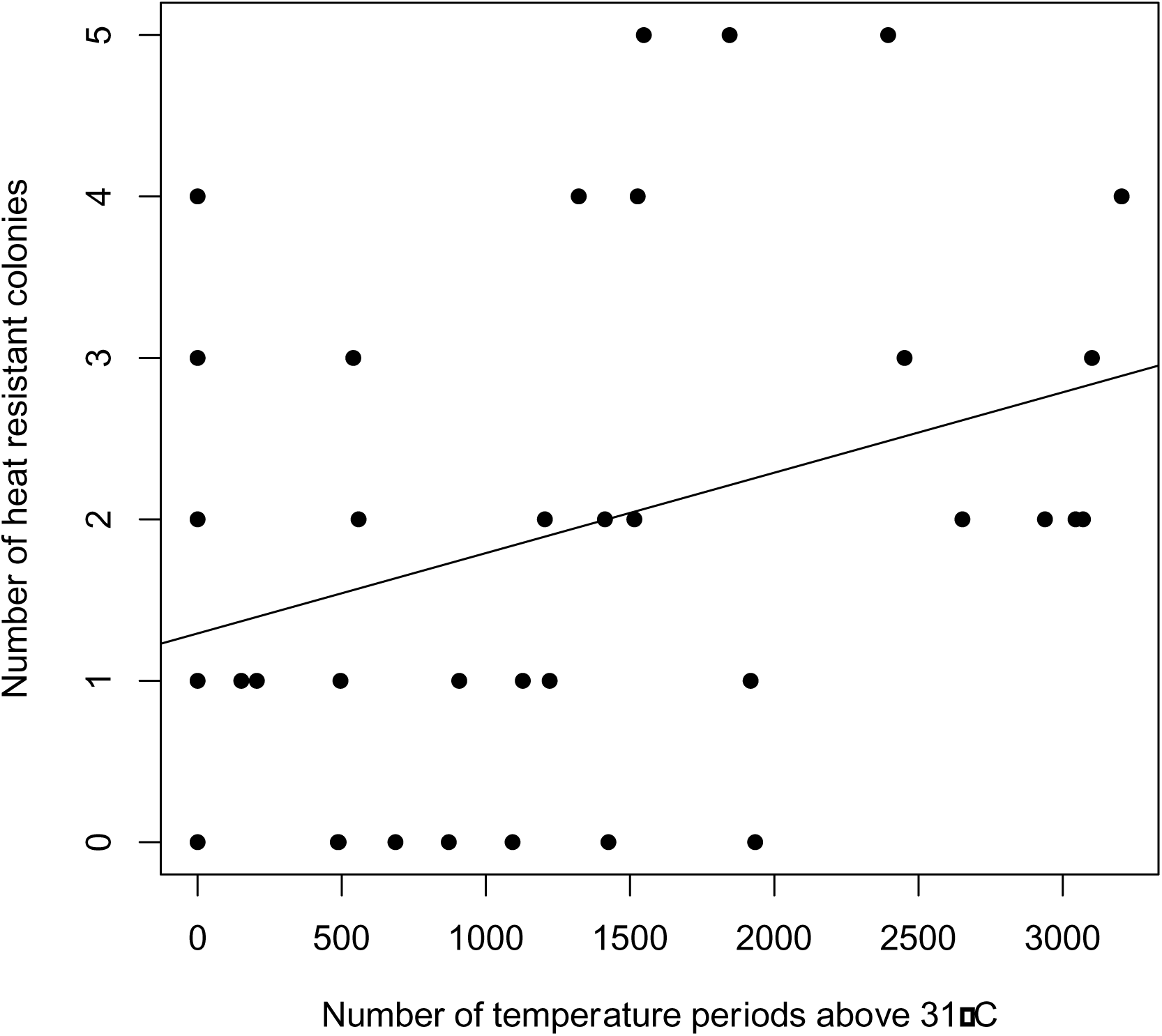
The relationship between reef temperature and the number of bleaching resistant individuals. Correlation of heat resistant colonies (number of colonies showing 2 of 3 criteria for high heat resistance) and temperature > 31°C -across reefs (R^2^ = 0.0922, p = 0.0338). Accompanying source data are available as Figure 4 – figure supplement 1 – Source Data.

**Figure 5 – figure supplement 1:**
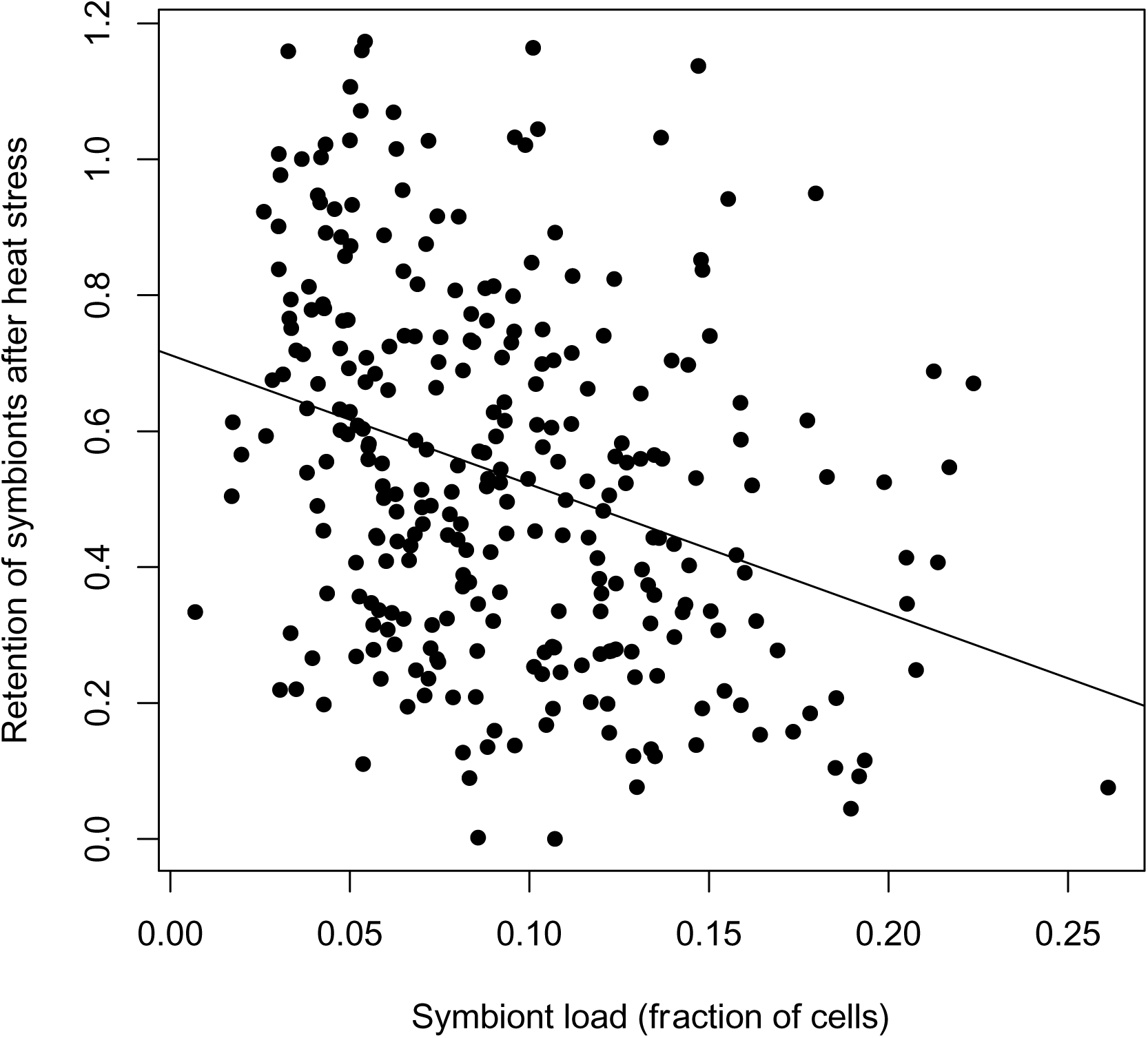
Symbiont load negatively correlates with symbiont loss. Corals with a higher proportion of symbiont cells in non-heated controls showed lower retention during heat exposure and higher overall bleaching. R^2^=0.1045, p=1.13×10^-8^. Accompanying source data are available as Figure 5 – figure supplement 1 – Source Data.

